# Impact of hepatitis E virus host- and virus-derived insertions on the antiviral response in primary human hepatocytes

**DOI:** 10.64898/2026.09.15.750637

**Authors:** Sarah Schlienkamp, Maximilian K. Nocke, Rainer G. Ulrich, Volker Kinast, Eike Steinmann, Daniel Todt

## Abstract

**Background & Aims:** Despite the global burden of hepatitis E virus (HEV) infection, host-virus interactions underlying HEV pathogenesis and progression to chronicity remain incompletely understood. This study aimed to characterize the host transcriptomic response to infection with HEV variants harboring different host- and virus-derived insertions in the hypervariable region (HVR), with a focus on innate immune activation.

**Methods:** We performed bulk RNA sequencing on PHHs from three donors infected with *in vivo-*identified HEV-3 variants harboring different HVR insertions (*RPS17*, HVR duplication, *SERPINA1*, *TRIM22*). To compare their replication fitness, we quantified progeny virus titers and viral RNA reads in the cellular transcriptome. Host transcriptomic responses were analyzed in detail at multiple time points to assess virus-induced differential gene expression. To further characterize the response to HEV infection with different HVR insertion variants, we performed gene ontology analysis and identified individual HEV-induced genes.

**Results:** Progeny virus titers at 72 hours post infection (h p.i.) were comparable across all PHH donors and HVR insertion variants. Two donors mounted robust host responses to HEV infection, characterized by widespread gene deregulation and interferon stimulated gene (ISG) upregulation at 72 h p.i. The HVR duplication insertion variant consistently triggered a weaker host response compared to *RPS17*, *SERPINA1* and *TRIM22* variants, with significantly fewer differentially expressed genes and reduced ISG induction. Gene ontology term analysis revealed enrichment of antiviral pathways upon infection with the different insertion variants, which was less pronounced for the HVR duplication variant.

**Conclusions:** These findings reveal that HEV HVR rearrangement variants elicit distinct host response patterns.Importantly, our results suggest a positive correlation of viral replication and host response induction. This indicates that the HVR may play a crucial role in shaping host-virus interactions.

**Impact and implications:** This study provides novel insights into HEV-host interactions, characterizing donor-specific and variant-dependent differences in antiviral responses. Our findings highlight the importance of understanding the role of the HVR in chronic HEV infection, a major clinical challenge particularly in immunocompromised patients where persistent viremia can lead to progressive liver damage. The observed variations in innate immune activation patterns across different donors and viral variants suggest a critical role of innate immunity in shaping HEV pathogenesis and potentially determining infection outcomes.

## Introduction

Hepatitis E virus (HEV, *Paslahepevirus balayani*) accounts for 20 million human infections every year, making it a leading cause for acute viral hepatitis and a significant global health burden [1]. A recent World Health Organization (WHO) update estimated 3.3 million symptomatic infections annually and reported 4,431 deaths caused by HEV infection in 2023 [2]. Although self-limiting in most cases, HEV infection can result in fulminant hepatitis and liver failure, particularly in vulnerable populations such as pregnant women and immunocompromised individuals, who are generally at elevated risk of developing severe illness upon HEV infection [3, 4]. In immunocompromised patients like solid organ transplant recipients, HEV infection can additionally progress to chronicity, which often complicates antiviral therapy through the emergence of treatment resistance mutations [5–7]. Treatment options remain limited to the off-label use of ribavirin (RBV), a broad-spectrum antiviral, and pegylated interferon-alpha (peg-IFNα), both of which are associated with severe side effects [3]. A deeper understanding of HEV-host interactions is therefore essential to broaden the knowledge on drivers of severe and chronic infection and to enable the development of effective antiviral therapies for prevention and treatment.

HEV is organized within the family *Hepeviridae*, with the species *Paslahepevirus balayani* being the major agent causing human HEV infections, while exhibiting a broad host tropism that varies by genotype [8, 9]. Human infections are typically caused by *Paslahepevirus balayani* genotypes 1-4 (HEV-1-4), among which HEV-1 and HEV-2 are strictly anthroponotic with a waterborne transmission route, while HEV-3 and HEV-4 are zoonotic with pigs serving as the main natural reservoir and source of transmission to humans. The 7.2 kilobase (kb) HEV genome consists of positive-sense single-stranded RNA ((+)ssRNA) and comprises three open reading frames (ORF1-3). Viral replication depends primarily on the nonstructural polyprotein encoded by ORF1, which includes essential enzymes such as a helicase (Hel) and the RNA-dependent RNA polymerase (RdRp). Beyond its role in replication, ORF1 also modulates the host’s immune response to create favorable conditions for viral replication [10, 11]. Within ORF1, the hypervariable region (HVR) - a poorly conserved, proline-rich domain - has recently been identified to underlie the intra-host genomic variability mainly reported in chronically infected patients. This variability arises from partial insertions of host-derived sequences as well as duplications of viral genomic regions [8, 12–19]. The role of HVR insertions in modulating viral fitness has further been demonstrated in the context of the first HEV strains cultured [20, 21]. Collectively, these findings suggest that modifications within the HVR shape virus-host interactions, particularly regarding the regulation of host antiviral defenses [22]. In this study, we employed primary human hepatocytes (PHHs) to investigate how previously identified host- and virus-derived HVR insertions affect the host innate immune response under physiologically relevant conditions.

## Material and Methods

### Plasmids

The plasmid encoding the full-length viral genome of the Kernow-C1/p6 virus isolate (HEV-3; GenBank accession Nr.: JQ679013) was a gift from Suzanne U. Emerson. Further plasmids carrying alternative HVR insertions are based on this backbone where the Kernow-C1/p6 *RPS17* insertion was substituted with respective gene sippets, generated via PCR-based mutagenesis, overlap-extension PCR and Gibson Assembly, as published previously [18].

### Cell Culture

Human hepatoma HepG2 cells (American Tissue Culture Collection, ATCC, Nr.: HB-8065) were grown in Dulbeccós Modified Eaglés Medium (DMEM-high-glucose, Gibco, #11965) supplemented with 10% (v/v) fetal calf serum (FCS, Capricorn, Lot. Nr. CPC21-4114), 1% (v/v) non-essential amino acids (NEAAs, Gibco, #11140050), 2 mM L-glutamine (Gibco, #25030), 100 IU/mL penicillin and 100 µg/mL streptomycin (Gibco, #15140) (DMEM complete). HepG2/C3A cells (kindly provided by Charles Rice, Rockefeller University, USA) were grown in Minimum Essential Medium (MEM, Gibco, #11095) supplemented with 10% (v/v) ultra-low immunoglobulin G (IgG) FCS (Gibco, #16250-078), 1% (v/v) NEAAs, 2 mM L-glutamine, 100 µg/mL gentamicin (Gibco, #15710) and 1 mM sodium pyruvate (Gibco, #11360). HepG2 and HepG2/C3A cells were grown on cell culture dishes (Sarstedt) coated with rat collagen (SERVA Electrophoresis, #47256.01). Cryopreserved PHHs (donors #CHM2420, #HH240404 and #HH250124, see Table S1) were obtained from Primacyt (Schwerin, Germany). Thawing and seeding on collagen-coated 24-well plates was performed following the distributor’s instructions. For cultivation, Human Hepatocyte Maintenance Medium (HHMM, Primacyt) was used. All cells were incubated at 37 °C at 5% (v/v) CO_2_.

### Production of cell culture-derived HEV (HEV_CC_)

Infectious HEV_CC_ particle production was performed as previously described [23]. Briefly, HepG2 cells were transfected via electroporation with in vitro-transcribed HEV Kernow-C1/p6 RNA. Intracellular non-enveloped HEV_CC_ was prepared from cell lysates by resuspension of the cells in DMEM seven days post transfection (d.p.t.), followed by three freeze–thaw cycles using liquid nitrogen and ice. Cell debris was removed by centrifugation at 10,000 × g for 10 min. The clarified supernatant was aliquoted and stored at -80 °C until usage. Virus titers were determined by titration onto HepG2/C3A cells and fixation followed by ORF2 capsid protein staining seven days post infection (d.p.i.) to quantify focus-forming units per mL (FFU/mL) via immunofluorescence microscopy.

### HEV infection of PHHs

For HEV infection, PHHs were thawed and seeded following the distributor’s instructions. The next day, cells were inoculated at a multiplicity of infection (MOI) of 0.2 with HEV Kernow-C1/p6-derived variants carrying the different HVR insertions. The next day, the culture medium was exchanged with fresh HHMM and the cells were washed thrice with phosphate-buffered saline (PBS) to remove remaining inoculum. At 8, 12, 24, 48 and 72 hours post infection (h p.i.) the cells were washed thrice with PBS, lysed and total RNA was extracted using the NucleoSpin RNA kit (Macherey-Nagel, 740955.50) according to the manufacturer’s instructions. In addition, viral titers were determined at 72 h p.i. by washing thrice with PBS, harvesting of intracellular progeny virus and retitration on HepG2/C3A cells as described above.

### Immunofluorescence staining and microscopy

For immunofluorescence staining, cells were fixed with 3% paraformaldehyde (PFA, Roth, 93351) for 15 min, followed by three PBS washes and permeabilization with 0.2% Triton X-100 (Roth, 3051.3) in PBS for 5 min. Cells were washed thrice again with PBS and blocked with 5% horse serum (HS) in PBS for at least 1 h at room temperature (RT) on a rocking shaker. Polyclonal HEV-3 (ORF2)-specific rabbit hyperimmune serum [24] was applied at 1:5,000 dilution in 5% HS and incubated overnight at 4 °C under permanent agitation. Cells were then washed thrice and incubated with a secondary antibody (donkey anti-rabbit AlexaFluor488, Invitrogen, A-21206; 1:1,000 in 5% HS) for 2-3 h at RT on a rocking shaker. Cells were washed again three times prior to staining of the nuclei with 4′,6-diamidino-2-phenylindole (DAPI, Invitrogen, D1306; 1:10,000 in H_₂_O) and three more washing steps with PBS. Stained cells were stored in PBS until imaging. Fluorescence images were acquired with a wide-field fluorescence microscope (Keyence BZ-X800E) using a 4 × 0.75-numerical aperture (NA) air-objective. DAPI (358 nm) and ORF2 capsid protein (488 nm) signals were acquired sequentially by using the BZ-X Filter DAPI and BZ-X Filter GFP, respectively.

### RNA sequencing

Further processing and Illumina sequencing of total RNA extracted from PHHs was performed in-house. Quality of RNA was assessed by NanoDrop One (Thermo Scientific) and the 4150 TapeStation (Agilent) to determine the RNA Integrating Number (RIN). Sequencing libraries were generated using polyA enrichment and libraries were sequenced on an Illumina NextSeq 1000 in 50-mer single-read mode (SR50) with an anticipated sequencing depth of 20 million reads per sample (50-mer paired-end mode (PE50) for one sample: donor 3, *TRIM22*, 48 h p.i.). RNAseq data have been deposited in National Center for Biotechnology Information’s Gene Expression Omnibus (GEO) and are accessible through GEO Series accession number GSE347085 (https://www.ncbi.nlm.nih.gov/geo/query/acc.cgi?acc=GSE347085).

### Bioinformatic analysis

A pipeline script to coordinate processing the raw sequencing data to raw gene counts and transcripts per million (TPM) was written in Julia (version 1.12.7) [25]. Raw sequencing data was trimmed by sequencing quality and for sequencing adapters utilizing Trimmomatic (version 0.40) [26]. Sequence mapping was first performed against viral side calling Tanoti (https://github.com/vbsreenu/Tanoti) with specified genomes matching each construct. Functions to correct false positive mapping of human reads to insertions within the viral references were implemented in Julia employing the packages FASTX (version 2.1.7) (https://github.com/BioJulia/FASTX.jl) and XAM (version 0.4.2) (https://github.com/BioJulia/XAM.jl). All unmapped and corrected reads were provided to STAR (version 2.7.11b) [27] alongside the human reference (GRCh38.p14). The mapped data was filtered and sorted by utilizing SAMtools (version 1.21) [28], to eliminate optical duplicates, Picard MarkDuplicates (version 3.1.1) (https://broadinstitute.github.io/picard/) was called in SAMtools output. Genes were counted using featureCounts (version 2.1.1) [29], while TPM calculation was implemented in Julia.

Principal component analysis (PCA), differentially expressed gene (DEG) and gene ontology (GO) analyses were conducted in R (version 4.6.1) [30]. The PCA was performed on TPM values. In order to define a suitable TPM cutoff for DEG analyses, we identified the intersection point of the TPM density curves for non-coding DNA sequence (CDS) and CDS genes from all samples. This resulted in a TPM cutoff of 2.465115, while the fold change (FC) threshold was set to be 3. edgeR (version 4.10.3) [31] was utilized to perform DEG analyses for all timepoints within donor 1 and 2. FC tables for both donors’ 72 h samples were filtered by our defined cutoffs. Intersections between the resulting genes were computed. The clusterProfiler (version 4.20.0) [32] function enrichGo was called on these intersections to execute a gene set enrichment analysis per construct. GO terms were reduced by rrvgo (version 1.24.0) [33]. Data visualization was performed using the packages ggplot2 (version 4.0.3) [34], pheatmap (version 1.0.13) [35] and ComplexUpset (version 1.3.3) [36].

### Image processing software

Images were analyzed with ImageJ version 1.54f (https://imagej.net/software/fiji/). Graphs were plotted with GraphPad Prism version 11.0.2 for Windows (https://www.graphpad.com). Final graphics were edited using Adobe Illustrator 2024 (https://www.adobe.com/).

## Results

### HEV infection elicits HVR insertion variant-specific replication kinetics in PHHs

To elucidate the transcriptional host response induced by HEV variants harboring distinct HVR insertions, we conducted total RNA sequencing of both mock-infected and HEV-infected PHHs. In our experimental setup, we included the frequently used Kernow-C1/p6 strain naturally containing the *RPS17* HVR insertion (further referred to *RPS17*), alongside three additional variants harboring HVR insertions derived from host genes and viral sequences. These variants were previously identified in a patient undergoing RBV treatment failure and have been demonstrated to enhance HEV replication *in vitro* to an extent comparable to the *RPS17* insertion [18]. Specifically, we included the host-derived insertions *SERPINA1*.1 and h.*TRIM22* (further referred to as *SERPINA1* and *TRIM22*) and dup1, a duplication of an HVR snippet (further referred to as dup) (Figure 1A). To mitigate potential confounding effects through immune response-triggering compounds present in the inoculum, we employed conditioned medium derived from HepG2/C3A cells as mock inoculum for uninfected control cells, which was prepared in the same manner as the infectious virus inoculum. At 72 h p.i. we assessed the generation of infectious progeny virus from HEV-inoculated PHHs by harvesting intracellular virus and subsequent retitration onto HepG2/C3A cells (Fig. 1B and C). In donor 1, viral titers spanned from 9.5×10^4^ FFU/mL (dup variant) to 3.7×10^5^ FFU/mL (*SERPINA1*), representing the highest values observed across all donors. Donor 2 exhibited comparatively lower titers between 1.8×10^4^ FFU/mL (*RPS17*) and 4×10^4^ FFU/mL (*TRIM22*). While donors 1 and 2 displayed only modest variations in progeny virus titers across the different HVR insertion variants, donor 3 exhibited the most pronounced variations. Specifically, we observed comparable titers for *TRIM22* and *RPS17* (3.2×10^5^ FFU/mL and 3.6×10^5^ FFU/mL, respectively), a 3-fold increase for *SERPINA1* (9.6×10^5^ FFU/mL) and most notably a 133-fold reduction for the dup variant (2.4×10^3^ FFU/mL), relative to *RPS17*.

**Figure 1:**
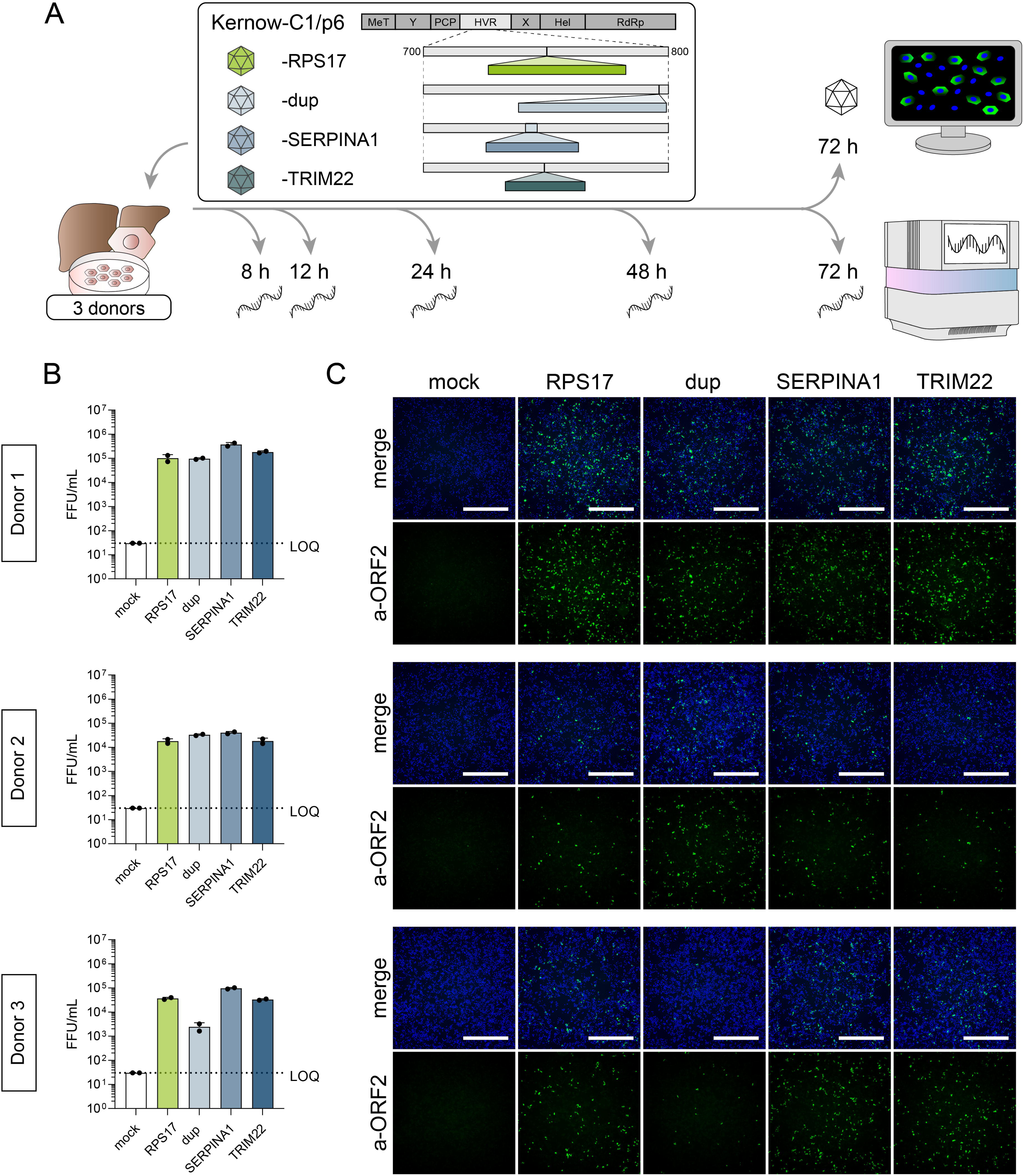
Quantification of intracellular progeny virus reveals robust infection of primary human hepatocytes (PHHs) with HVR insertion variants. (**A**) PHHs of 3 donors were infected with HEV variants harboring HVR insertion from host gene snippets (*RPS17*, *SERPINA1*, *TRIM22*) or the viral genome (HVR sequence duplication, dup), which are all drawn to scale. Conditioned medium served as mock infection control. Total RNA was extracted from PHHs at 8, 12, 24, 48 and 72 h p.i. and supplied to Illumina sequencing. In addition, intracellular infectious progeny virus particles were harvested at 72 h p.i. and quantified by retitration and immunofluorescence microscopy. (**B**) Quantification of progeny virus from 3 individual PHH donors infected with HEV HVR insertion variants. Viral particles were harvested at 72 h p.i. and retitrated on HepG2/C3A cells in duplicates (dots). Titers are depicted as means +standard deviation (SD) of the calculated focus forming units per mL (FFU/mL). Dashed line represents the limit of quantification (LOQ). (**C)** Representative immunofluorescence images of progeny virus retitration onto HepG2/C3a cells. Viral capsid protein (ORF2) = green; DAPI = blue. Scale bars = 1 mm. Abbreviations: PHHs, primary human hepatocytes; MeT, methyltransferase; Y, Y-domain; PCP, papain-like cysteine protease; HVR, hypervariable region; X, X-domain; Hel, helicase; RdRp, RNA-dependent RNA polymerase; h p.i., hours post infection; LOQ, limit of quantification.

To further evaluate viral replication kinetics, we aimed to compare replication fitness across donors and HVR insertion variants by quantifying temporal changes in HEV-specific reads within the cellular transcriptome. RNA samples were collected from both mock-infected and HEV-infected PHHs at 8, 12, 24, 48 and 72 h p.i., thereby capturing early steps of the replication cycle including the completion of viral entry and later stages of established infection (Fig. 1A). Since some of the used HEV constructs contain sequence insertions derived from the host genome, we developed a correction method to ensure accurate transcript assignment to either the viral or the host genome (Fig. 2A). This novel approach prevents false positive mappings caused by HVR insertions and avoids biases in our quantitative transcriptome analysis. After validating consistent coverage of our mapped data over the HEV genome (Fig. 2B), we quantified HEV replication kinetics (Fig. 2C). This analysis revealed an increase of HEV replication after 24 h p.i. compared to the 8 hour baseline for all HVR insertion variants with the exception of dup in donor 3. Despite slight peaks at 48 h p.i. observed for some insertion variants in donors 1 and 2, the percentage of mapped viral reads relative to host reads at 72 h p.i. remained in the range of up to 1% across all variants. Thus, the temporal replication profiles at 72 h p.i. reflect our progeny virus titration data, demonstrating only slight variations in viral replication profiles across the HVR insertion variants. Yet, the dup variant lacking peaks at 48 h p.i. in all donors displayed altered replication kinetics. Similarly, peaks at 48 h p.i. could not be detected in donor 3, suggesting that replication efficiency at this time point was generally lower.

**Figure 2:**
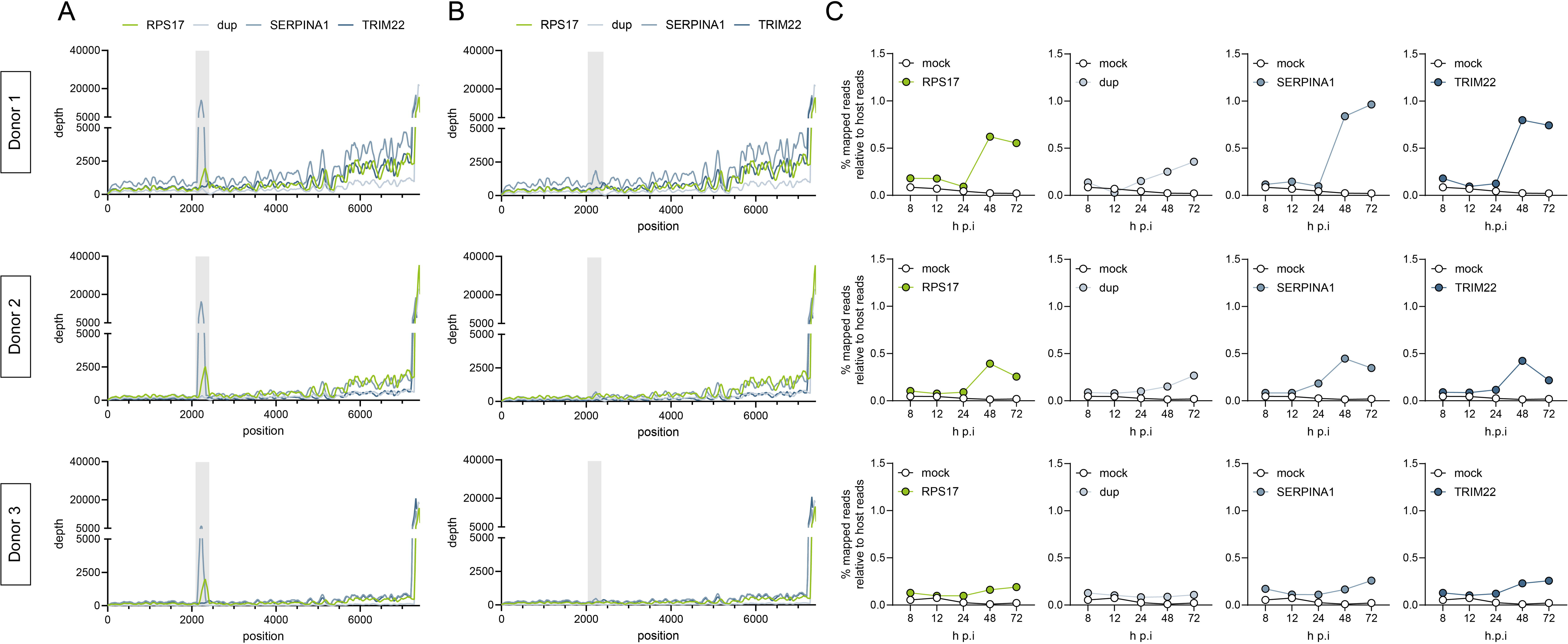
RNA-sequencing of viral reads reveal robust replication of HVR variants in PHHs. **(A)** Coverage of mapped reads along the 7.2 kb HEV genome before analysis pipeline adaptation. Individual plots represent data for all four HVR insertion variants per donor. Areas highlighted in gray represent the HVR. **(B)** Coverage of mapped reads along the HEV genome after removal of host-derived insertion bias. Individual plots represent data for all four HVR insertion variants per donor. Areas highlighted in gray represent the HVR. **(C)** Percentage of mapped HEV reads relative to reads mapped to the host over the course of the experiment. Rows represent insertion variants per individual donor. Abbreviations: HVR, hypervariable region; h p.i., hours post infection.

### RNA-sequencing reveals virus-induced differential gene expression at 72 hours post infection in PHHs

To confirm the integrity of the PHHs over the time of the experiment, we analyzed the expression patterns of cellular marker genes as well as host factors involved in HEV infection and a subsequent antiviral response (Fig. 3A). At both 8 h p.i. and 72 h p.i., pulmonary and neuronal markers were expressed at low levels, whereas hepatic markers were strongly expressed, confirming the stability and authenticity of the PHHs up to the final time point and excluding substantial dedifferentiation in culture over time. Moreover, the robust expression of host factors associated with HEV entry, replication and egress [37–48], along with interferon (IFN) receptors, Pattern Recognition Receptors (PRRs) and immune-related transcription factors provided a suitable model to study both HEV infection and the infection-induced host response.

**Figure 3:**
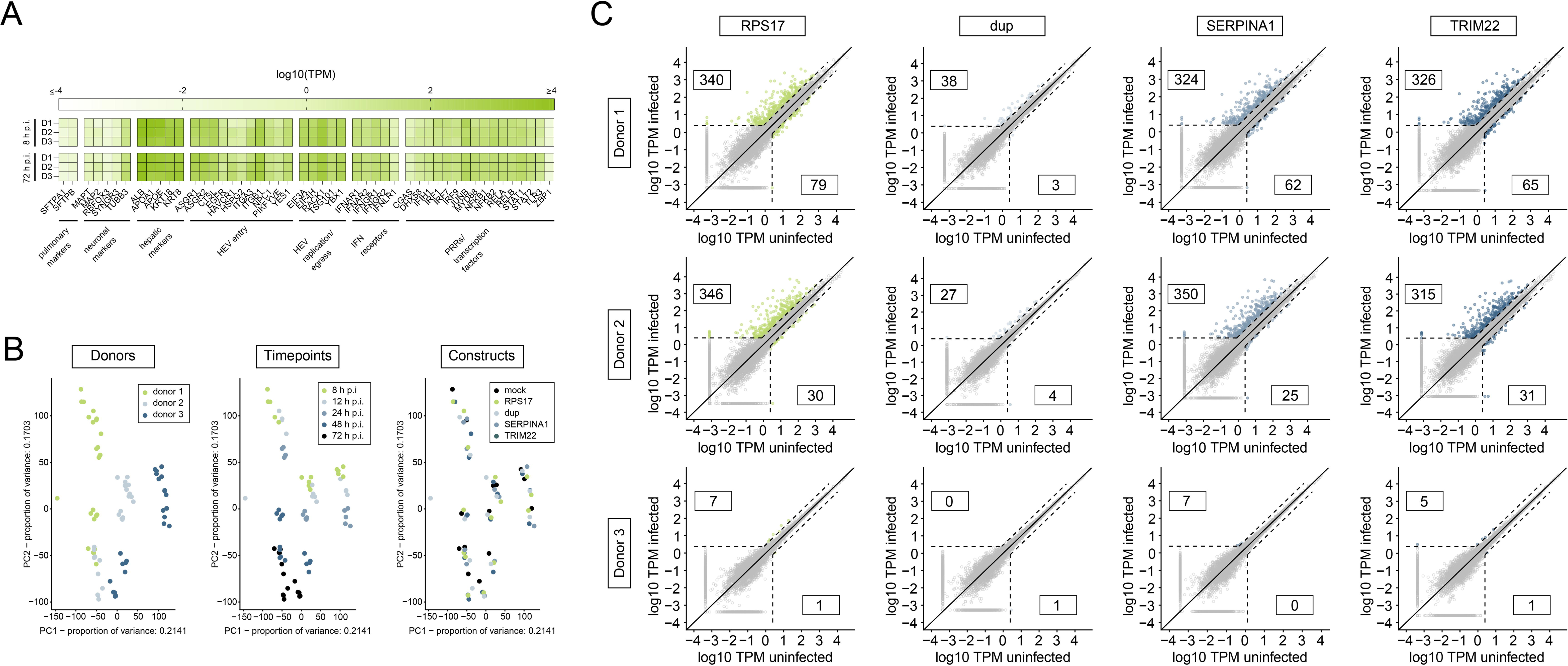
RNA-sequencing reveals virus-induced differential host gene expression at 72 hours post infection in PHHs. **(A)** Expression of tissue-specific marker genes (pulmonary, neuronal and hepatic), HEV host factors (entry, replication, egress), interferon receptors, pattern recognition receptors and transcription factors in the 3 individual PHH donors. Depicted are TPM expression data from mock-infected samples at 8 and 72 h p.i. **(B)** Principal component analysis of transcriptome data grouped by donor, time point and HVR insertion variant. **(C)** Differentially expressed genes in infected PHH samples (y-axis) at 72 h p.i. compared to the respective mock-infected control (x-axis). Rows represent differential expression upon infection with HVR insertion variants per individual donor. Dashed lines depict cutoffs of at least 3-fold change in expression and 2.47 TPM before or after infection. Included counts of DEGs above the cutoffs (colored dots) represent robustly up- or downregulated genes. Abbreviations: TPM, transcripts per million; IFN, interferon; PRRs, pattern-recognition receptors; PHH, primary human hepatocyte; PC, principal component; HVR, hypervariable region; h p.i., hours post infection; DEGs, differentially expressed genes.

Next, we analyzed the host response upon infection with the different HEV HVR insertion variants. To assess global changes in the host transcriptome upon HEV infection, we performed a PCA on the RNA-sequencing data. PCA was employed to reduce dimensionality while presenting the highest share of data variance, enabling us to identify patterns based on donors, time points and HVR insertions (Fig. 3B). Samples segregated into defined clusters based on donor identity, underscoring a natural donor-to-donor variability in the transcriptional profiles. Over the course of our experiment, samples displayed comparable time-dependent trajectories reflecting similar gene expression alterations from the earliest time point to the latest. In contrast, PCA did not reveal distinct clustering between mock and infected samples or the different HVR insertion variants, indicating that inoculum-dependent differences in the host transcriptome are either subtle or masked by donor-specific and temporal effects. We further identified donor 1 at 12 h p.i. with the dup variant as outlier among our RNA-sequencing samples (Fig. S1) and therefore excluded this sample from further analyses. Together, these data suggest time-dependent transcriptomic changes in the three donors, that are mostly unrelated to infection status.

To examine the impact of HEV infection on gene expression for the individual donors and HVR insertion variants, we compared the transcriptome of infected PHHs to the corresponding mock-infected control of the respective time point. We filtered for robustly differentially expressed genes (DEGs) at the latest time point by defining a cutoff of 2.47 TPM and a 3-fold change over the mock control (Fig. 3C, Fig. S2). In donors 1 and 2, we observed a profound modulation of host gene expression at 72 h p.i. in response to *RPS17*, *SERPINA1* and *TRIM22* variants, with approximately 300-350 upregulated and 20-80 downregulated genes (Fig. 3C). In contrast, the dup variant elicited only a mild response to infection, inducing less than 40 upregulated and less than 5 downregulated genes. In donor 3, the number of DEGs was markedly lower across all insertion variants compared to the other two donors, with less than 10 upregulated genes and a maximum of 1 downregulated gene. Of note, these host response patterns resembled our viral replication dynamics data, suggesting that the extent of the host gene dysregulation upon HEV infection positively correlates with viral replication fitness.

To systematically assess shared and unique host transcriptional responses, we generated UpSet plots for the final time point under two comparative frameworks [49]. First, insertion-wise comparisons within individual donors revealed substantial overlap in DEGs among *RPS17*, *SERPINA1*, and *TRIM22* variants, with markedly reduced overlap when the dup variant was included (Fig. 4A). Second, donor-wise comparisons within each insertion variant revealed that donors 1 and 2 displayed highly similar expression patterns across *RPS17*, *SERPINA1*, and *TRIM22* variants, whereas donor 3 showed minimal deregulation (Fig. 4B). As also demonstrated previously in our DEG analysis, the dup variant consistently exhibited considerably fewer deregulated genes across all donors, with no DEGs shared between all 3 donors. Collectively, these findings indicate that donors 1 and 2 mount robust and largely overlapping host responses to HEV infection, while donor 3 shows minimal transcriptional changes, suggesting a less pronounced host response in this donor. To detect potential discrepancies in the innate immune activation between insertion variants and donors, we quantified IFN expression as surrogate marker of antiviral signaling, early after infection and at the final time point (Fig. 4C). In both infected and mock samples, we observed low expression of especially *IFNK* and type III IFNs at 8 h p.i., suggesting a cellular stress response to plating rather than an early antiviral response. At 72 h p.i., we detected elevated levels of IFNB1 and type III IFNs exclusively in infected samples, confirming an active innate immune response to HEV infection. Notably, the reduced number of deregulated genes previously observed for the dup variant and donor 3 corresponded with diminished IFN expression upon infection. Importantly, in both cases, a less pronounced overall host response correlates with lower replication rates compared to other insertion variants and donors, respectively.

**Figure 4:**
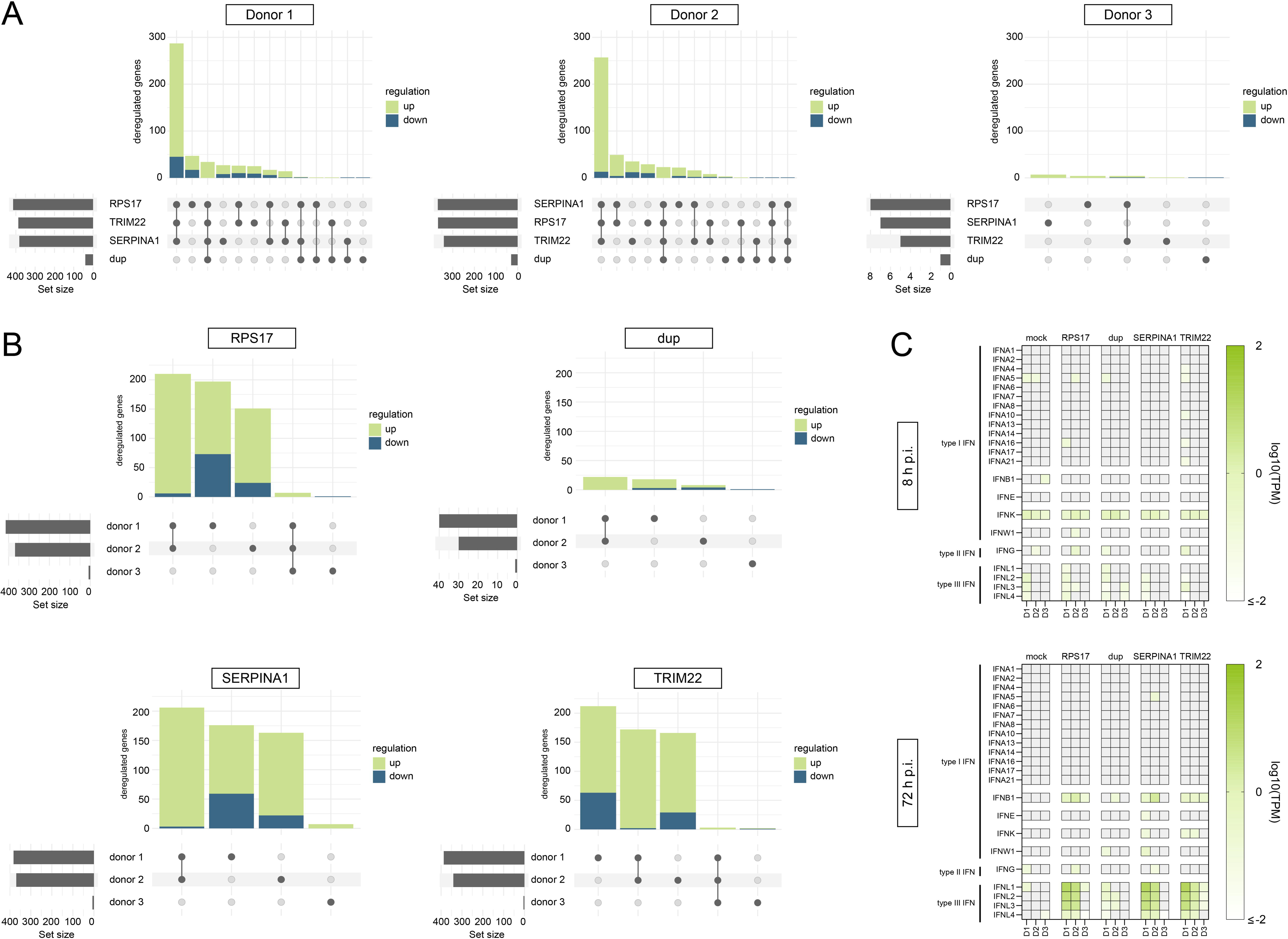
PHHs from responsive donors 1 and 2 elicit comparable antiviral responses. **(A)** Number of DEGs per donor for each construct at 72 h p.i. and their unique and intersecting gene counts. The gray bar plots show the number of DEGs per insertion variant. Bar plots on top represent the amount of unique and intersecting DEGs per insertion variant or insertion variant set. **(B)** Number of DEGs per construct for each donor at 72 h p.i. and their unique and intersecting gene counts. **(C)** Interferon expression at 8 (top) and 72 (bottom) h p.i. with the different HVR insertion variants. Depicted are log_10_ TPM values detected in the 3 individual PHH donors. Gray color indicates TPM values of 0. Abbreviations: DEGs, differentially expressed genes; h p.i., hours post infection; HVR, hypervariable region; TPM, transcripts per million; PHH, primary human hepatocytes.

### Virus-induced differential gene expression and affected gene ontology terms are shared between PHH donors

Focusing our further transcriptomic analysis on the virus-induced innate immune response, we excluded donor 3 due to the lack of a robust IFN expression. To mask their heterogenous genetic background, we compared individual DEGs upon HEV infection by statistical analysis of pooled data from both donors. Specifically, we generated MA-plots to highlight DEGs exceeding our previously determined cutoff threshold for each donor separately and for genes shared between both donors (Fig. 5A). Our analysis confirmed that the majority of DEGs above the cutoff were shared between donors 1 and 2 and robustly deregulated across both donors. Additionally, we observed that the dup variant triggered minimal activation of host response pathways, consistent with our previous findings.

**Figure 5:**
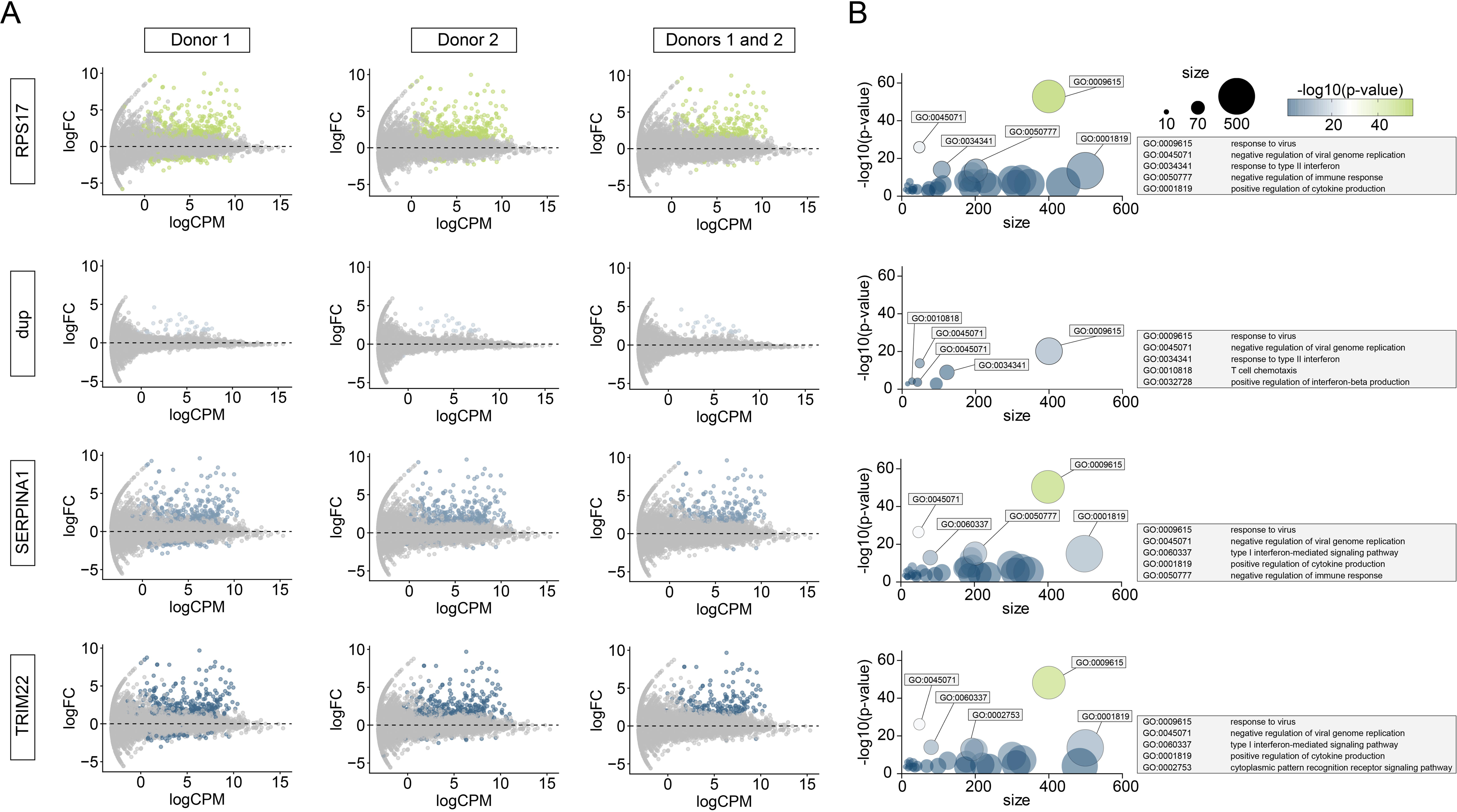
Responsive PHH donors 1 and 2 share virus-induced differential gene expression and affected gene ontology terms. **(A)** MA plots visualizing the gene regulation for donor 1 and 2 in each HVR insertion variant at 72 h p.i. Highlighted DEGs in the first two columns are corresponding to the ones identified per donor in Fig. 3. The third column represents the intersection of those DEGs. **(B)** GO term analyses of DEG intersections at 72 h p.i. per HVR insertion variant. The circle size corresponds to the number of genes per GO term. The top 5 GO terms ranked by score are labeled. Abbreviations: PHH, primary human hepatocyte; h p.i., hours post infection; DEGs, differentially expressed genes; GO, gene ontology.

We further processed the genes above our defined cutoffs in a GO analysis for donors 1 and 2, to compare the regulation of cellular pathways at 72 h p.i. with the different insertion variants (Fig. 5B). For all insertion variants, we found the same top two regulated GO terms “response to virus” and “negative regulation of viral genome replication”. Further GO terms related to antiviral defense mechanisms were shared among the insertion variants including “response to type II interferon”, “negative regulation of immune response” and “positive regulation of cytokine production”. Overall, all top regulated GO terms across the insertion variants stand in the context of viral infection and host response. However, we observed that the number of regulated GO terms was strongly decreased for the dup variant compared to the other insertions. To analyze this finding in more detail, we characterized the host response upon infection by identifying strongly dysregulated individual genes upon infection, for which we used the lists of genes above our predefined cutoffs at 72 h p.i., again (Fig. S3). Upon infection with the variants *RPS17*, *SERPINA1* and *TRIM22* we observed a downregulation of genes connected to general cellular metabolism (*HMGCS2*, *ADH1B* and *ALDOB*, among others). Among infection-induced genes we mainly identified ISGs, IFNs, cytokines and other genes related to an innate immune response, with *CCL5* representing the strongest upregulation across all insertion variants. Although a number of classical ISGs such as *OASL*, *OAS2*, *OAS3* and *MX1* were induced by all insertion variants, the number of regulated genes upon infection was dramatically lower for the dup variant. Similarly, we observed a strong induction of core-ISGs, a set of ISGs identified by Shaw et al. to be highly conserved among mammalian species [50], upon infection with the *RPS17*, *SERPINA1* and *TRIM22* variant at 72 h p.i., but not for the dup variant (Fig. S4).

## Discussion

Due to its suspected role as critical modulator of virus-host interactions, the HVR within the ORF1-encoded nonstructural polyprotein of HEV has recently gained attention as a potential determinant of viral pathogenesis and host tropism. Host-derived insertions and viral sequence duplications within this region are associated with intra-host variability and tissue culture adaptation. In recent years, multiple genomic rearrangements of the HVR have been identified mainly in immunosuppressed patients with chronic HEV infection, many of which have been demonstrated to enhance viral replication in tissue culture [17–19]. Together with the typical rapid elimination of non-beneficial virus variants among populations of small RNA viruses, this suggests a key role of the HVR sequence heterogeneity in viral fitness. However, the mechanisms underlying the insertion of sequences as well as their specific consequences regarding virus-host interactions need to be identified yet. Moreover, the precise impact of distinct HVR insertion variants on antiviral host responses remains unclear. In our study we addressed this gap by providing a comprehensive and authentic characterization of the host transcriptomic response to HEV infection, for which we infected PHHs with multiple *in vivo* identified HVR insertion variants (*RPS17*, dup, *SERPINA1* and *TRIM22*) in a standardized HEV genome backbone (Kernow-C1/p6) [18].

Our data revealed that two out of three donors mount robust antiviral responses to HEV infection, characterized by widespread gene deregulation and ISG induction at 72 h p.i.. In contrast, the third donor exhibited minimal transcriptional changes, suggesting that, despite the inoculation of the donors with a normalized MOI, viral replication levels in this donor were not meeting the threshold to trigger antiviral pathways. This donor-specific variability aligns with previous reports of inter-individual differences in antiviral capacity and susceptibility to virus infection, *in vitro* and *in vivo* as well as for HEV and other RNA viruses [5, 51, 52]. The phenomenon of donor-specific variability can also be observed *in vivo* as discrepancies in clinical outcomes of HEV-infections and a higher susceptibility of certain individuals to chronic HEV infection.

Despite achieving comparable progeny virus titers at 72 h p.i. to other variants, the dup variant exhibited slightly distinct replication kinetics. While *RPS17*, *SERPINA1* and *TRIM22* variants showed peaks in viral reads at 48 h p.i., the dup variant lacked this peak. This specific replication pattern potentially alters direct or indirect interactions of the dup insertion variant with host immune pathways compared to other variants. In line with this, our analysis revealed a less pronounced induction of IFNB1 and type III IFNs at 72 h p.i. with the dup variant. Given that HEV initially suppresses early IFN responses [15, 53], which is suspected to only permit antiviral gene activation once replication reaches a critical threshold, the altered IFN induction of the dup variant might reflect a viral strategy to balance replication and immune evasion, as previously proposed for other hepatotropic viruses [54, 55]. In our further transcriptomic comparison of the HVR insertion variants, the dup variant consistently triggered a markedly weaker host response across all donors, with significantly fewer DEGs and reduced ISG induction compared to *RPS17*, *SERPINA1* and *TRIM22*. This aligns with growing evidence that HEV may indirectly regulate host responses via HVR variations by modulating replication fitness, though the exact mechanisms remain unclear [18, 22]. In line with this, lower replication levels of the dup variant compared to the other HVR insertion variants might contribute to the less pronounced antiviral response observed.

In this context, our findings complement and expand upon those of Paronetto et al., who recently reported a stronger host immune response upon infection with HEV insertion variants of higher replication fitness [22]. While their study employed cancer cell lines, we utilized PHHs. Yet, our results demonstrate a similar correlation between replication fitness and host immune response, particularly observed in the gene regulation patterns for the dup variant that align with altered replication kinetics. Notably, whereas Paronetto et al. included a host-derived insertion variant that does not enhance replication fitness, our analysis evaluated the impact of a virus-derived insertion variant (dup) known to enhance viral replication compared to the Kernow-C1/p1 strain lacking the *RPS17* insertion, thereby providing additional insights into how different types of HVR insertions may differentially modulate host-virus interactions [18].

In addition to viral replication fitness, the unique nature of the dup insertion, being derived from viral sequences rather than host-derived insertions like *RPS17*, *SERPINA1*, and *TRIM22*, could explain its phenotype. Host-derived insertions might mimic cellular proteins more efficiently compared to viral proteins and thereby modulate host pathways through molecular mimicry, potentially enabling more efficient adaptation. In contrast, the ORF1-derived dup insertion from viral origin may lack these pre-adapted interactions with host immune machinery, resulting in a different interaction profile with the host antiviral responses. This could explain why the dup variant elicits a reduced host response activation. Of note, previous studies have discovered distinct effects of host- and virus-derived insertions on the proportions of negatively and positively charged amino acid residues [14, 56]. However, while both increases and decreases in negatively charged residues have been reported across different studies, the overall direction and functional consequences of these changes remains unclear to date.

In conclusion, our study demonstrates that HEV HVR rearrangement variants elicit distinct host response patterns, with the dup insertion variant exhibiting less prominent innate immune activation patterns compared to the other insertion variants. These findings underscore the potential role of the HVR in modulating host-virus interactions and highlight the need for systematic HVR characterization to elucidate its contribution to HEV pathogenesis.

## List of Abbreviations

HEV: hepatitis E virus
HVR: hypervariable region
PHH: primary human hepatocyte
ISG: interferon-regulated gene
GO: gene ontology
RBV: ribavirin
MOI: multiplicity of infection
PRR: pattern-recognition receptor
PCA: principle component analysis
DEG: differentially expressed gene
IFN: interferon
FC: fold change
TPM: transcripts per million
FFU: focus forming units
h p.i.: hours post infection

## Acknowledgments

We are grateful to Suzanne Emerson for the HEV Kernow-C1/p6 clone. We thank all members of the Department for Molecular and Medical Virology, Ruhr-University Bochum, and the Department of Translational and Computational Infection Research (TRACiR), Ruhr-University Bochum, for helpful support, suggestions, and discussions.

## Declaration of generative AI and AI-assisted technologies in the manuscript preparation process

During the preparation of this work, the authors used Mistral Small 4 119B 2603 to refine language and grammar. The authors reviewed and edited the output as needed and take full responsibility for the content of the published article.

## Authors contributions

Performed research: S.S., M.K.N.

Analysed data: S.S., M.K.N.

Designed research: S.S., V.K., E.S., D.T.

Provided resources: R.G.U.

Visualized data: S.S., M.K.N.

Supervised the project: V.K., E.S., D.T.

Wrote the original draft: S.S.

All authors provided critical feedback and contributed to shaping the research, analysis, and manuscript.

---

**Figure S1:** Clustering of normalized gene expression values of complete dataset. Presented values are z-scores of transcripts per million (TPM) clustered by rows and columns.

**Figure S2:** Virus-induced differential gene expression from 8 to 48 hours post infection. Differentially expressed genes in infected (y-axis) PHH samples compared to the respective mock-infected control (x-axis) at 8 h p.i. (A), 12 h p.i. (B), 24 h p.i. (C) and 48 h p.i. (D). Rows represent differential expression upon infection with HVR insertion variants per individual donor. Dashed lines depict cutoffs of at least 3-fold change in expression and 2.47 TPM before or after infection. Included counts of DEGs above the cutoffs (colored dots) represent robustly up- or downregulated genes. Abbreviations: PHH, primary human hepatocytes; h p.i., hours post infection; HVR, hypervariable region; TPM, transcripts per million; DEGs, differentially expressed genes.

**Figure S3:** Gene regulation of differentially expressed gene intersections between donor 1 and 2 for each HVR insertion variant. Log10 TPMs for infected and uninfected samples at 72 h p.i. are presented next to their corresponding log2 FCs. Heatmap values are ordered by decreasing log2 FC. Abbreviations: TPM, transcripts per million; FC, fold change.

**Figure S4:** Core-ISG induction upon infection with HVR insertion variants. Heatmap with detailed deregulation of individual core-ISGs [1] in PHHs from three independent donors (labeled below) at 8 and 72 h p.i. with different HVR insertion variants, quantified as log2 FC over the corresponding mock-infected sample. Abbreviations: ISG, interferon-stimulated gene; HVR, hypervariable region; FC, fold change; PHHs, primary human hepatocytes; h p.i., hours post infection.

**Table S1:**
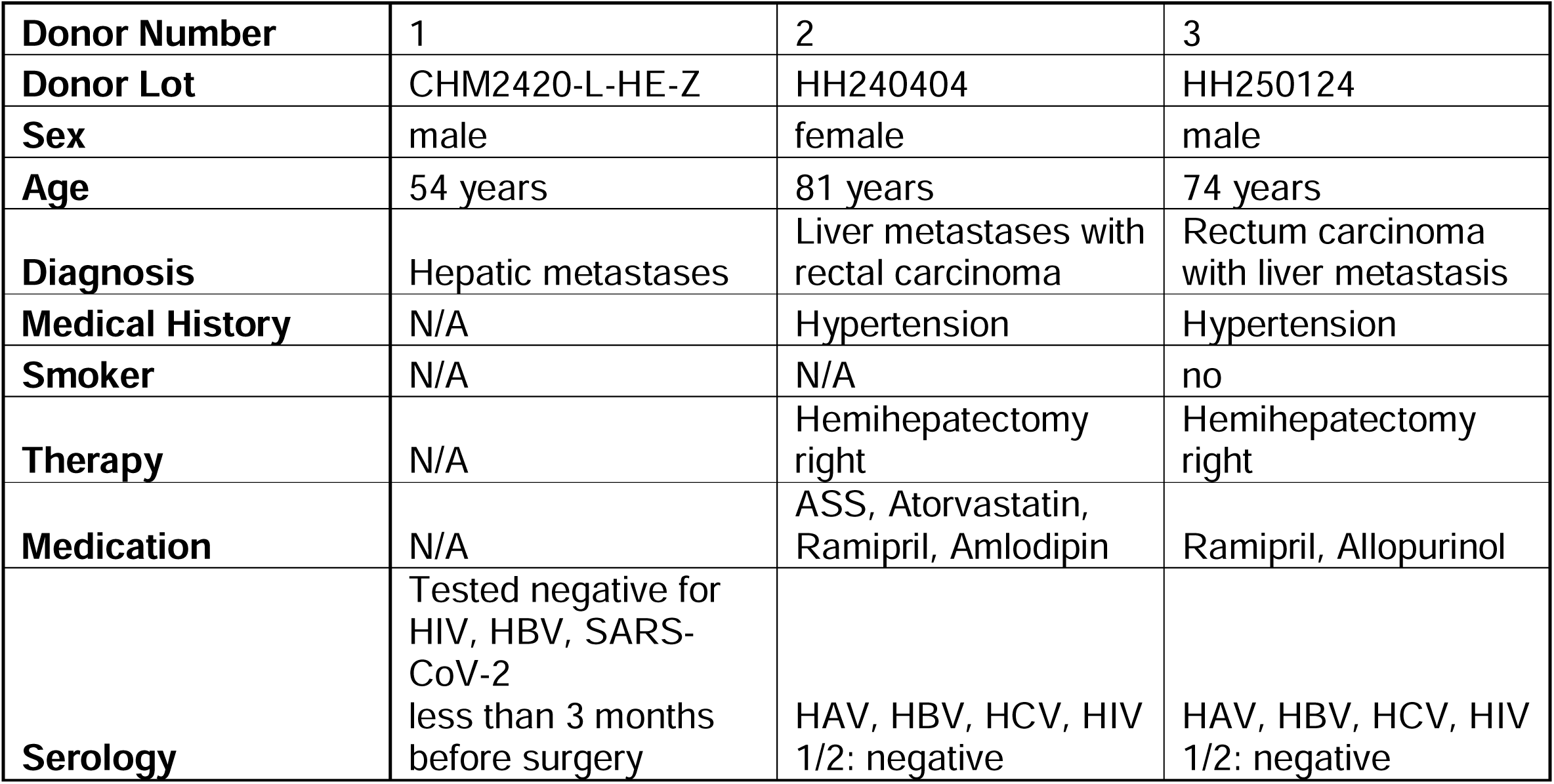
Donor information of primary human hepatocytes used in this study. Abbreviations: N/A, not available; HAV, hepatitis A virus; HBV, hepatitis B virus; HCV, hepatitis C virus; HIV, human immunodeficiency virus; SARS-CoV-2, severe acute respiratory syndrome coronavirus 2.

