## Supplement for "Impact of hepatitis E virus host- and virus-derived insertions on the antiviral response in primary human hepatocytes"

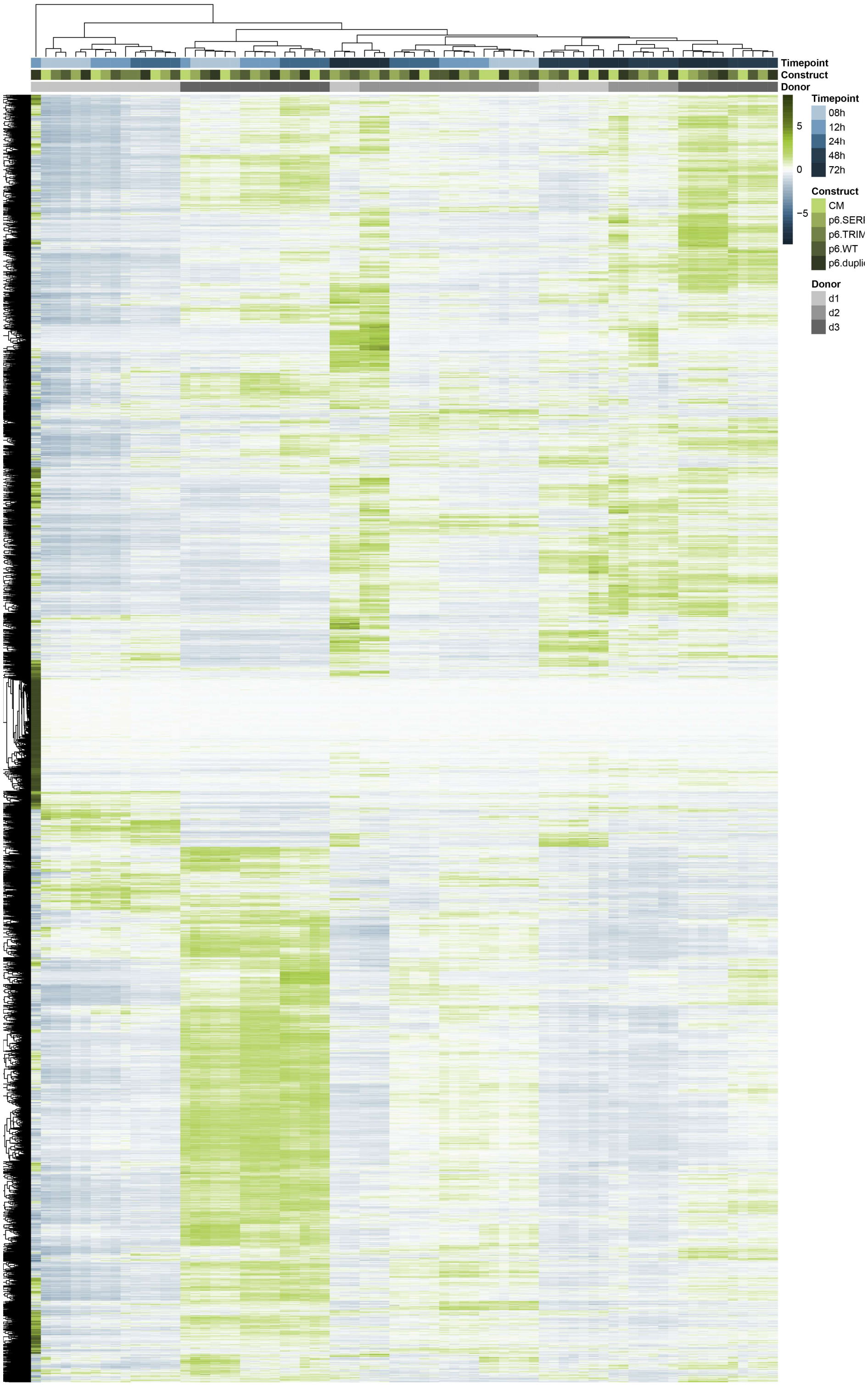


**Figure S1: Clustering of normalized gene expression values of complete dataset.** Presented values are z-scores of transcripts per million (TPM) clustered by rows and columns.

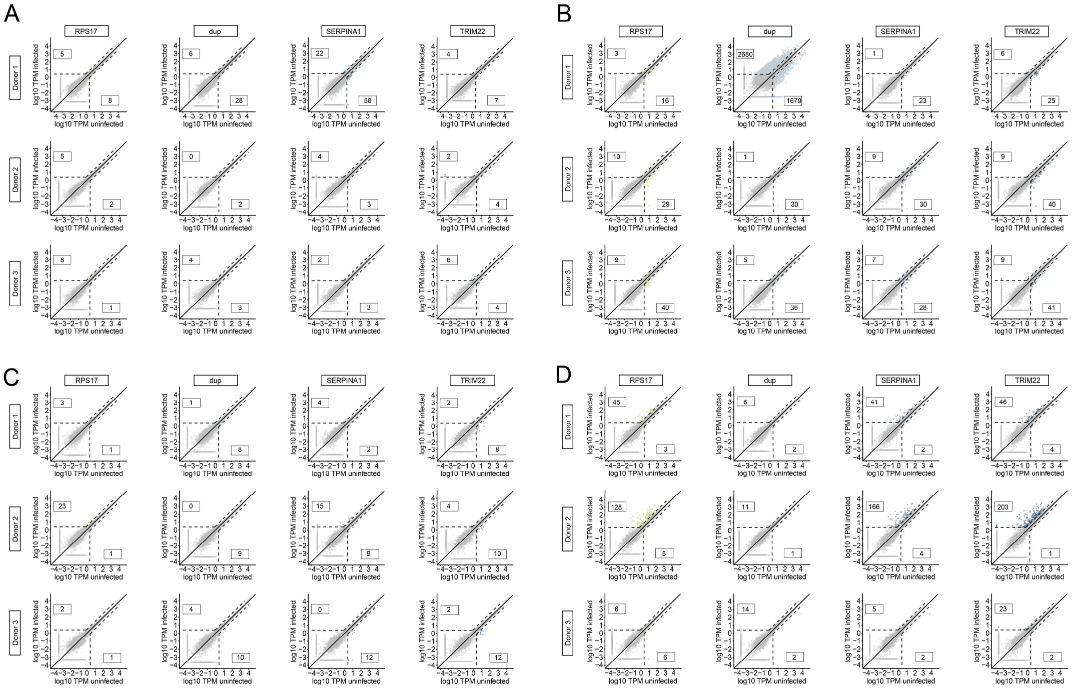


**Figure S2: Virus-induced differential gene expression from 8 to 48 hours post infection.** Differentially expressed genes in infected (y-axis) PHH samples compared to the respective mock-infected control (x-axis) at 8 h p.i. **(A)**, 12 h p.i. **(B)**, 24 h p.i. **(C)** and 48 h p.i. **(D)**. Rows represent differential expression upon infection with HVR insertion variants per individual donor. Dashed lines depict cutoffs of at least 3-fold change in expression and 2.47 TPM before or after infection. Included counts of DEGs above the cutoffs (colored dots) represent robustly up- or downregulated genes. Abbreviations: PHH, primary human hepatocytes; h p.i., hours post infection; HVR, hypervariable region; TPM, transcripts per million; DEGs, differentially expressed genes.

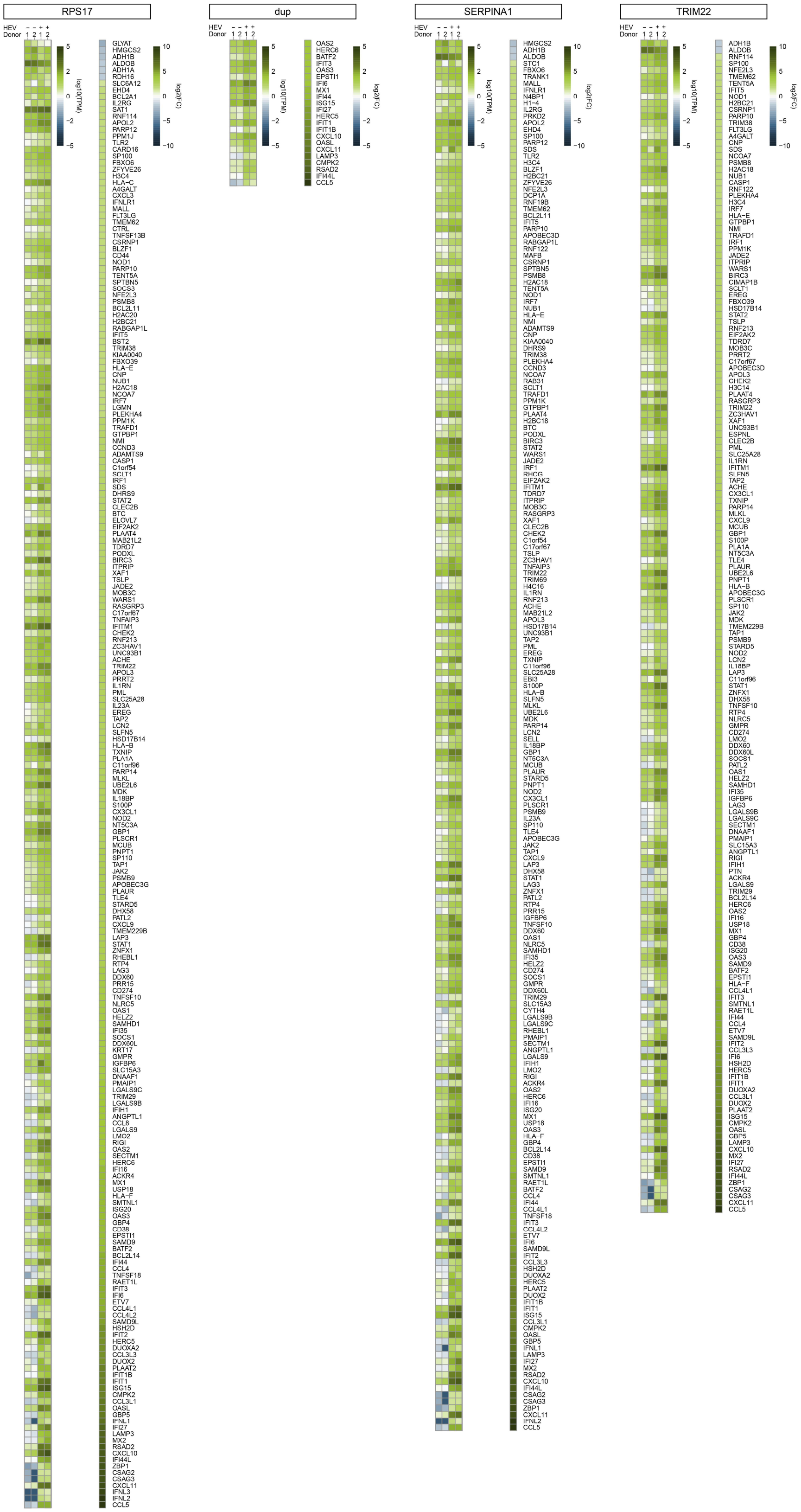
RPS17
dup
SERPINA1
TRIM22
HEV
Donor
log10(TPM)
log2(FC)

**Figure S3: Gene regulation of differentially expressed gene intersections between donor 1 and 2 for each HVR insertion variant.** $Log_{10}$ TPMs for infected and uninfected samples at 72 h p.i. are presented next to their corresponding $log_2$ FCs. Heatmap values are ordered by decreasing $log_2$ FC. Abbreviations: TPM, transcripts per million; FC, fold change.

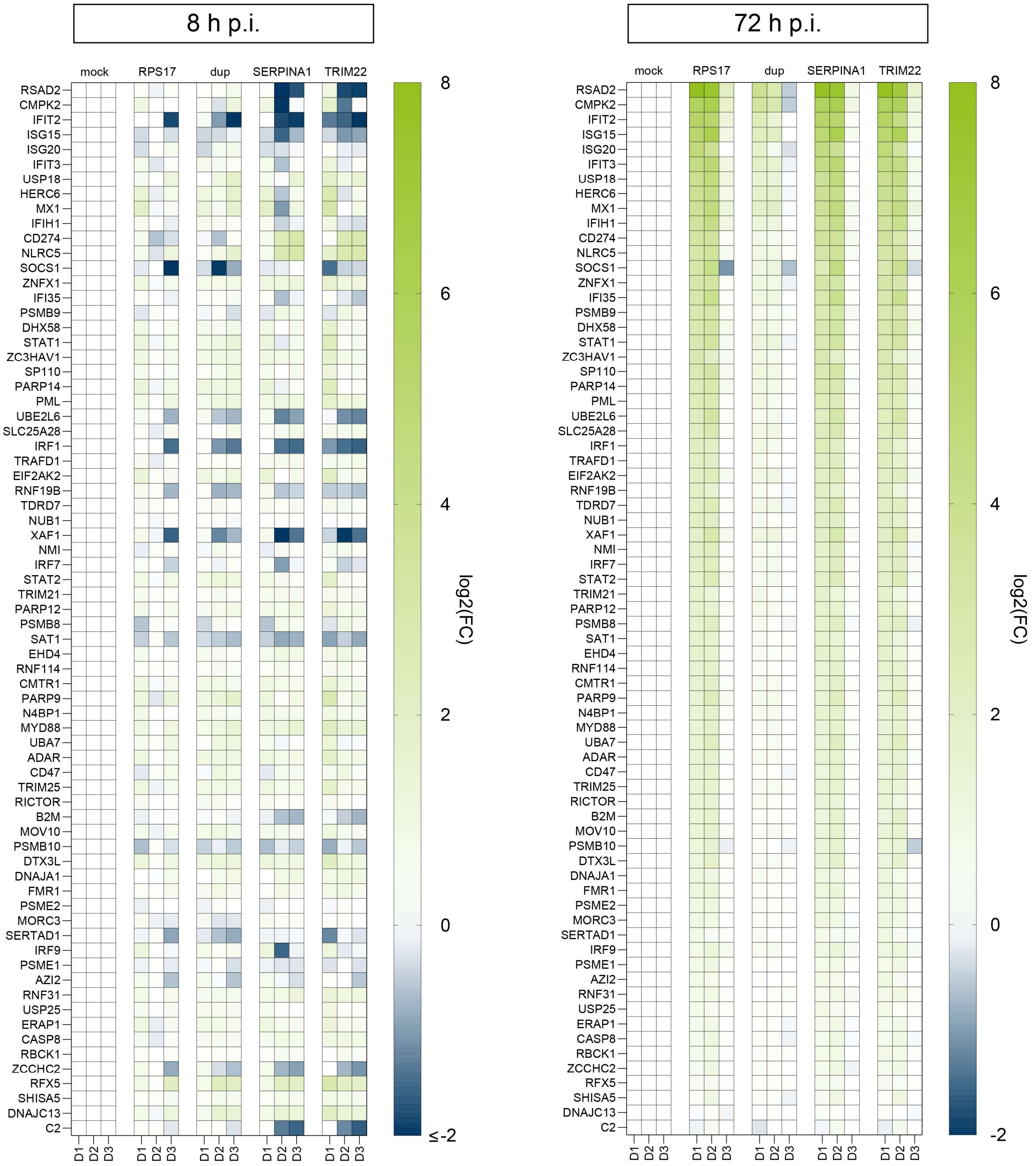


**Figure S4: Core-ISG induction upon infection with HVR insertion variants.** Heatmap with detailed deregulation of individual core-ISGs [1] in PHHs from three independent donors (labeled below) at 8 and 72 h p.i. with different HVR insertion variants, quantified as $log_2$ FC over the corresponding mock-infected sample. Abbreviations: ISG, interferon-stimulated gene; HVR, hypervariable region; FC, fold change; PHHs, primary human hepatocytes; h p.i., hours post infection.

**Supplementary tables:**

**Table S1: Donor information of primary human hepatocytes used in this study.** Abbreviations: N/A, not available; HAV, hepatitis A virus; HBV, hepatitis B virus; HCV, hepatitis C virus; HIV, human immunodeficiency virus; SARS-CoV-2, severe acute respiratory syndrome coronavirus 2.

| **Donor Number** | 1 | 2 | 3 |
|---|---|---|---|
| **Donor Lot** | CHM2420-L-HE-Z | HH240404 | HH250124 |
| **Sex** | male | female | male |
| **Age** | 54 years | 81 years | 74 years |
| **Diagnosis** | Hepatic metastases | Liver metastases with rectal carcinoma | Rectum carcinoma with liver metastasis |
| **Medical History** | N/A | Hypertension | Hypertension |
| **Smoker** | N/A | N/A | no |
| **Therapy** | N/A | Hemihepatectomy right | Hemihepatectomy right |
| **Medication** | N/A | ASS, Atorvastatin, Ramipril, Amlodipin | Ramipril, Allopurinol |
| **Serology** | Tested negative for HIV, HBV, SARS-CoV-2 less than 3 months before surgery | HAV, HBV, HCV, HIV 1/2: negative | HAV, HBV, HCV, HIV 1/2: negative |
